## Supplementary File 1 for "Evolutionary and ecological processes influencing chemical defense variation in an aposematic and mimetic *Heliconius* butterfly"

**Table S1A.** Cyanogenic glucoside compound concentrations (mean, standard deviation (SD) and *P*-values of Tukey HSD pairwise comparisons) in natural populations of *Heliconius erato* and *Passiflora biflora* sampled along environmental gradients in Panama and Ecuador. Statistically significant pairwise comparisons are marked in bold.

***H. erato*** — Linamarin %

| Country | Population | Mean | SD | Pairwise comparisons (Tukey HSD *P*-value) *Dry* | *Int* | *Wet* | *Low* | *High* |
|---|---|---|---|---|---|---|---|---|
| Panama | *Dry* | 0.771 | 0.557 | | | | | |
| Panama | *Intermediate* | 0.776 | 0.465 | 0.997 | | | | |
| Panama | *Wet* | 0.569 | 0.356 | 0.445 | 0.295 | | | |
| Ecuador | *Low* | 0.011 | 0.028 | **<0.001** | **<0.001** | **<0.001** | | |
| Ecuador | *High* | 0.026 | 0.033 | **<0.001** | **<0.001** | **<0.001** | 0.911 | |

***H. erato*** — Lotaustralin %

| Country | Population | Mean | SD | Pairwise comparisons (Tukey HSD *P*-value) *Dry* | *Int* | *Wet* | *Low* | *High* |
|---|---|---|---|---|---|---|---|---|
| Panama | *Dry* | 0.293 | 0.177 | | | | | |
| Panama | *Intermediate* | 0.495 | 0.218 | **<0.001** | | | | |
| Panama | *Wet* | 0.337 | 0.198 | 0.766 | **0.007** | | | |
| Ecuador | *Low* | 0.000 | 0.000 | **<0.001** | **<0.001** | **<0.001** | | |
| Ecuador | *High* | 0.000 | 0.000 | **<0.001** | **<0.001** | **<0.001** | 1.000 | |

***H. erato*** — Total cyanogen %

| Country | Population | Mean | SD | Pairwise comparisons (Tukey HSD *P*-value) *Dry* | *Int* | *Wet* | *Low* | *High* |
|---|---|---|---|---|---|---|---|---|
| Panama | *Dry* | 1.064 | 0.709 | | | | | |
| Panama | *Intermediate* | 1.271 | 0.631 | 0.432 | | | | |
| Panama | *Wet* | 0.907 | 0.521 | 0.905 | 0.098 | | | |
| Ecuador | *Low* | 0.011 | 0.028 | **<0.001** | **<0.001** | **<0.001** | | |
| Ecuador | *High* | 0.026 | 0.033 | **<0.001** | **<0.001** | **<0.001** | 0.936 | |

***P. biflora***

| Country | Population | Passibiflorin (µg/mg) Mean | SD | Pairwise comparisons (Tukey HSD *P*-value) *Dry* | *Int* | *Wet* | Passibiflorin triglycoside (µg/mg) Mean | SD | Pairwise comparisons (Tukey HSD *P*-value) *Dry* | *Int* | *Wet* |
|---|---|---|---|---|---|---|---|---|---|---|---|
| Panama | *Dry* | 11.005 | 9.459 | | | | 0.732 | 0.620 | | | |
| Panama | *Intermediate* | 20.291 | 12.925 | **0.035** | | | 1.166 | 0.654 | 0.126 | | |
| Panama | *Wet* | 17.095 | 7.528 | 0.123 | 0.653 | | 0.911 | 0.497 | 0.598 | 0.478 | |

| Country | Population | Tetraphyllin A (µg/mg) Mean | SD | Pairwise comparisons (Tukey HSD *P*-value) *Dry* | *Int* | *Wet* | Total cyanogens (µg/mg) Mean | SD | Pairwise comparisons (Tukey HSD *P*-value) *Dry* | *Int* | *Wet* |
|---|---|---|---|---|---|---|---|---|---|---|---|
| Panama | *Dry* | 0.113 | 0.416 | | | | 11.850 | 10.073 | | | |
| Panama | *Intermediate* | 0.087 | 0.289 | 0.970 | | | 21.544 | 13.586 | **0.038** | | |
| Panama | *Wet* | 0.000 | 0.000 | 0.448 | 0.709 | | 18.005 | 7.953 | 0.146 | 0.627 | |

**Table S1B.** Variance components of *Heliconius* biosynthesized cyanogenic toxicity estimated with REML animal models. For each variance component, Variance (V), Standard error (*S.E.*) of variance, Statistical significance (*P*) of variance, and Proportion of variance of total phenotypic variance ($V/V_P$) are shown. In addition, broad-sense heritability $H^2$ ($V_G/V_P$) and evolvability $e_\mu$ ($V_G/\mu^2$; % ) are calculated based on the genetic variance component (shaded columns). Data consists of 322 observations, including 20 mothers, 13 fathers, and 289 F1 offspring belonging to 20 broods (of which 18 are full-siblings, and 2 are paternal half-siblings).

| | | | Genetic variance component, heritability and evolvability | | | | | Maternal variance | | | | Fixed effects | | | | | | | | Residual variance | | | Total phenotypic variance |
|---|---|---|---|---|---|---|---|---|---|---|---|---|---|---|---|---|---|---|---|---|---|---|---|
| | | | | | | | | | | | | Feeding treatment | | | | Sex | | | | | | | |
| Model type | Cyanogen trait | Trait mean ($\mu$) | $V_G$ | $V_G$ *S.E.* | *P* $V_G$ | $H^2$ ($V_G/V_P$) | $e_\mu$ ($V_G/\mu^2$); % | $V_{mat}$ | *S.E.* $V_{mat}$ | P $V_{mat}$ | $V_{mat}/V_P$ | $V_{trm}$ | *S.E.* $V_{trm}$ | $P_{trm}$ | $V_{trm}$ / $V_P$ | $V_{sex}$ | *S.E.* $V_{sex}$ | $P_{sex}$ | $V_{sex}/V_P$ | $V_R$ | *S.E.* $V_R$ | $V_R/V_P$ | $V_P$ |
| 1 | Total CNglc % | 0.772 | 0.0093 | 0.0069 | 0.0368 | 0.1152 | 1.5548 | | | | | 0.001 | 7.E-05 | 0.467 | 0.011 | 0.001 | 7.E-05 | 0.057 | 0.011 | 0.070 | 0.008 | 0.874 | 0.0805 |
| 1 | Linamarin % | 0.577 | 0.0063 | 0.0043 | 0.0202 | 0.1387 | 1.8869 | | | | | 5.E-05 | 4.E-06 | 0.418 | 0.001 | 0.001 | 5.E-05 | 0.036 | 0.013 | 0.038 | 0.004 | 0.848 | 0.0453 |
| 1 | Lotaustralin % | 0.195 | 4.0E-04 | 3.9E-04 | 0.1304 | 0.0668 | 1.0394 | | | | | 5.E-05 | 4.E-06 | 0.621 | 0.008 | 3.E-05 | 2.E-06 | 0.224 | 0.004 | 0.005 | 0.001 | 0.929 | 0.0059 |
| 2 | Total CNglc % | 0.772 | 1.0E-07 | 0.0312 | 1.0000 | 1.4E-06 | 1.7E-05 | 0.007 | 0.013 | 1.000 | 0.102 | 0.003 | 2.E-04 | 0.134 | 0.043 | 2.E-04 | 1.E-05 | 0.378 | 0.002 | 0.065 | 0.018 | 0.895 | 0.0727 |
