## Supplementary File 2 for "Evolutionary and ecological processes influencing chemical defense variation in an aposematic and mimetic *Heliconius* butterfly"

**Table S2:** Summary of *Heliconius erato* broods.

| | Mother id | Father id | Offspring n | Feeding treatment |
|---|---|---|---|---|
| | 1031 | N.A. | 7 | NO |
| | 1049 | 1029 | 6 | NO |
| | 1059 | 1037 | 7 | NO |
| | 1062 | N.A. | 12 | NO |
| | 1070 | N.A. | 10 | NO |
| | 1076 | 1030 | 10 | NO |
| | 1083 | 1030 | 9 | NO |
| | 1126 | N.A. | 9 | NO |
| | 1129 | 1100 | 8 | NO |
| | 1142 | N.A. | 7 | NO |
| | 1685 | N.A. | 12 | NO |
| | 1687 | 1421 | 8 | NO |
| | 1691 | 1657 | 6 | NO |
| | 1799 | 1664 | 5 | NO |
| | 1346 | 1291 | 38 | YES |
| | 1404 | 1238 | 42 | YES |
| | 1446 | 1224 | 20 | YES |
| | 1850 | 1722 | 25 | YES |
| | 1860 | 1729 | 16 | YES |
| | 2193 | 2120 | 32 | YES |
| **n** | **20** | **13** | **289** | |
| | | **Total n:** | **322** | |
