## Supplementary File 3 for "Evolutionary and ecological processes influencing chemical defense variation in an aposematic and mimetic *Heliconius* butterfly"

**SUPPLEMENTARY RESULTS**

**Reduced cyanogenic toxicity and increased nitrogen balance in drought-stressed *Passiflora* host plants**

Cyanogenic toxicity of the *Passiflora biflora* host plants varied environmentally, as demonstrated by significant effects of the full-factorial experimental treatments (Fig. S1, Table S3). Dry-treated plants (which received 1/3 of the water volume of wet-treated plants, mimicking conditions of drought) were less cyanogenic than were wet-treated plants ($F_{1, 95} = 13.2$, $P = 0.0004$; ANOVA of total CNglcs concentration including origin, treatment and their interaction). This effect was stronger in the plants originating from the Dry study area, with wet-treated plants being the most (mean = 8.72 µg/mg, sd = 2.62 µg/mg dry mass) and the dry-treated plants the less (mean: 5.03 µg/mg, sd: 3.62 µg/mg dry mass) toxic across all treatments (origin-by-treatment interaction $F_{1, 95} = 6.78$, $P = 0.011$). In contrast to the environmental effect, cyanogen concentration did not vary detectably among sites of origin ($F_{2, 95} = 0.53$, $P = 0.593$). The CNglcs chemical profile was similar across all plant treatment groups, with the major CNglcs compound passibiflorin constituting on average 95% of total CNglcs content, and the diglycoside of passibiflorin constituting the remaining proportion. An exception to this pattern occurred for plants from the Dry area under dry conditions (Dd treatment group), which were the only ones to contain a third type of CNglc, Tetraphyllin A, at up to 8% of the total CNglcs content (mean = 0.17 µg/mg dry mass, sd = 0.21 µg/mg dry mass).

In addition to cyanogen levels, plant quality indices representing nutritional values potentially important for *Heliconius* cyanogen biosynthesis varied across plant treatment groups (Fig. S1, Table S3). Most notably, the chlorophyll index increased ($F_{2, 310} = 19.4$, $P < 0.0001$) and the flavonoids index decreased ($F_{2, 309} = 21.8$, $P < 0.0001$) in the "dry"-treatment plants compared to "wet" and "st" treatment, which was reflected also in the increased nitrogen balance index (NBI) measuring nitrogen/carbon-ratio ($F_{2, 309} = 35.5$, $P < 0.0001$). This increase was especially strong in the plants originating from the "Dry" study area (origin-by-treatment interaction $F_{1, 309} = 9.4$, $P = 0.0023$). In contrast, the anthocyanin index increased in "wet"-treated plants ($F_{2, 304} = 77.5$, $P < 0.0001$), especially in the plants originating from the "Wet" study area (origin-by-treatment interaction $F_{1, 304} = 4.2$, $P = 0.0417$).

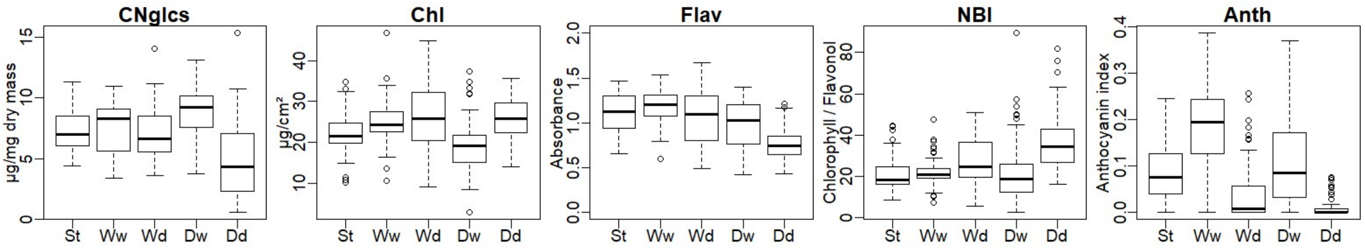


**Figure S1.** Cyanogen concentration and indices of plant quality in greenhouse-cultivated full-factorial treatment groups of *Passiflora biflora*, used as diet for laboratory broods of *Heliconius erato*. CNglcs: Total concentration of cyanogenic glucoside CNglcs compounds (µg/mg dry mass), Chl: chlorophyll content in µg/cm², Flav: epidermal flavonol content in absorbance units, NBI: nitrogen status, measured as the ratio between chlorophyll and flavonol and related to nitrogen/carbon allocation, Anth: anthocyanin index related to water-soluble plant pigments. *P. biflora* treatment groups are denoted as St; Standard cultivar, W/D; originates from "Wet"/"Dry" wild population along a rainfall gradient in Panama, respectively, w/d; cultivated in "wet"/"dry" conditions, respectively. See Table S3 for model results. Source data is deposited in Dryad data repository with DOI https://doi.org/10.5061/dryad.kprr4xh1w.

**Table S3.** Results of ANOVA models testing differences between host plant *Passiflora biflora* types cultivated in a full-factorial combination of plant origins and water availability treatments, and used as diet in common-garden rearing of *Heliconius erato*. The effects of watering treatment (dry vs. wet) and origin (Wet vs. Dry wild populations along a rainfall gradient in Panama) and their interaction were tested on *P. biflora* cyanogenic toxicity and other plant quality indices which represent nutritional values potentially important for *Heliconius* cyanogen biosynthesis. Plant traits: CNglcs; Total concentration of cyanogenic glucoside CNglcs compounds (µg/mg dry mass), Chl; chlorophyll content in µg/cm², Flav; epidermal flavonol content in absorbance units, NBI; nitrogen status, measured as the ratio between chlorophyll and flavonol and related to nitrogen/carbon allocation, Anth; anthocyanin index related to water-soluble plant pigments. Statistically significant (*P* < 0.05) results are highlighted in bold. Source data is deposited in Dryad data repository with DOI https://doi.org/10.5061/dryad.kprr4xh1w.

| | **Main model terms** | | | | | | | | | | | | **Post-hoc tests (TukeyHSD): *P-value*** | | | | | |
|---|---|---|---|---|---|---|---|---|---|---|---|---|---|---|---|---|---|---|
| | **Watering treatment** | | | | **Origin** | | | | **Treatment * Origin** | | | | **Watering treatment** | | | **Origin** | | |
| **Plant trait** | *F* | *D.F.* | *D.F. (res.)* | *P* | *F* | *D.F.* | *D.F. (res.)* | *P* | *F* | *D.F.* | *D.F. (res.)* | *P* | Dry - St | Wet - St | Wet - Dry | Dry - St | Wet - St | Wet - Dry |
| **CNglcs** | 13.25 | 1 | 95 | **4.4E-04** | 0.53 | 2 | 95 | 0.5926 | 6.78 | 1 | 95 | **1.1E-02** | 0.3022 | 0.3022 | **1.3E-03** | 0.7976 | 0.9844 | 0.5834 |
| **Chl** | 19.42 | 2 | 310 | **1.1E-08** | 23.57 | 1 | 310 | **1.9E-06** | 10.19 | 1 | 310 | **1.6E-03** | **5.6E-05** | 0.8149 | **0.0E+00** | 0.1715 | 0.0565 | **5.8E-06** |
| **Flav** | 21.82 | 2 | 309 | **1.4E-09** | 62.01 | 1 | 309 | **5.9E-14** | 5.23 | 1 | 309 | **2.3E-02** | **1.0E-07** | 0.6182 | **2.0E-07** | **1.1E-02** | **6.0E-04** | **0.0E+00** |
| **NBI** | 35.53 | 2 | 309 | **1.3E-14** | 9.15 | 1 | 309 | **2.7E-03** | 9.44 | 1 | 309 | **2.3E-03** | **0.0E+00** | 0.7707 | **0.0E+00** | 0.5041 | 0.3203 | **7.7E-03** |
| **Anth** | 77.47 | 2 | 304 | **2.0E-16** | 34.78 | 1 | 304 | **9.8E-09** | 4.18 | 1 | 304 | **4.2E-02** | **1.9E-06** | **5.0E-07** | **0.0E+00** | 0.0779 | **1.5E-02** | **0.0E+00** |
