## Supplementary File 4 for "Evolutionary and ecological processes influencing chemical defense variation in an aposematic and mimetic *Heliconius* butterfly"

**SUPPLEMENTARY METHODS**

### Further information on the study species *Heliconius erato*

*Heliconius erato* (Lepidoptera: Nymphalidae) is a widespread neotropical butterfly typically occurring at the edges of primary and secondary tropical rainforests. A striking diversity of color-pattern races with region-specific wing patterns have evolved as mimics sharing aposematic coloration with other species, typically *H. melpomene*, in order to deter common predators (Supple et al., 2013). *Heliconius erato* has a restricted home range and a dispersal distance of around 1-2 km per generation (Turner, 1971). The larvae can feed on several *Passiflora* species, depending on the local quantity and quality of available hosts (Kerpel & Moreira, 2005). In the Panamanian study area, the main natural hosts of *H. erato* are *P. biflora*, *P. coriaceae* and to a lesser extent *P. auriculata* (Merrill et al., 2013), while the preferred host plants in Ecuador are not well known (but possibly include *P. punctate*; (Hay-Roe & Nation, 2007). Cyanogenic toxicity of *H. erato* is moderate compared to other *Heliconius* species and arise from both biosynthesized and sequestered toxins, depending on the host on which it feeds (Arias et al., 2016; de Castro et al., 2019).

### Study sites and field collection of butterflies and host plants

*Heliconius erato demophoon* butterflies and its typical host plant *P. biflora* were collected from forest and forest-edge populations in Panama, and *H. erato lativitta* from the Eastern slope of the Andes in Ecuador. Butterflies were caught using a hand net and transported live to the laboratory. In Panama, sample collection took place in May (early wet season) 2017 within three study areas situated along a rainfall gradient from the Pacific to the Atlantic coast in the near vicinity of the Panama Canal in, including six different locations within the Dry, three locations in the Intermediate (Int), and along a 5 km forest road in the Wet study areas. Meteorological and hydrological data are collected in or in the immediate vicinity of the three study sites by the Smithsonian Tropical Research Institute (STRI) Physical Monitoring Program (https://biogeodb.stri.si.edu/physical_monitoring/). Although the distance between the two coasts is only about 65 km in the study region, it represents a steep rainfall gradient, with yearly rainfall (20-year average) of 3375 mm, 2720 mm, and 1881 mm in the Wet, Int, and Dry study areas, respectively. The average soil moisture content measured with a soil moisture meter (Digital Instruments Model PMS-714) at the time of sample collection was 27% (sd = 10%), 24% (sd = 10%), and 18% (sd = 8%) in the "Wet", "Int" and "Dry" study areas, respectively.

All individuals caught from the Panamanian study areas ($n$ = 92) were sexed and weighed, and males were preserved in 1 ml 100% MeOH on the day of capture. Before preservation as samples, females were kept in an outdoor insectary cage for 1-3 weeks (Dry, Int and Wet populations separately), where they were fed *ad libitum* with a 20% sugar solution enriched with protein concentrate supplemented with amino acids (Vetark Professional Critical Care Formula; two scoops per 1 l of sugar solution) and potted nectar plants (*Psychotria elata*), and were allowed to oviposit on potted *Passiflora biflora*. Of the butterfly samples, the body excluding wings and one half of the thorax were preserved in 1 ml 100% MeOH.

At the Ecuadorian sites,. At the Ecuadorian populations on the Eastern slope of the Andes, wild *H. erato lativitta* were collected in High (mean = 1200 m.a.s.l.) and Low altitude (mean = 400 m.a.s.l.). Fertilized females were brought to a common garden environment (insectaries at the University of IKIAM, Tena, Ecuador), allowed to oviposit and raised separately according to the altitude where they were collected and fed with *Passiflora punctata*. The thorax of 21 individuals from each altitude were preserved in 1 ml 100% MeOH.

*Passiflora* leaf samples were collected from the Panamanian Dry ($n$ = 20), Int ($n$ = 11) and Wet ($n$ = 20) study areas. The *Passiflora* collected from the Dry and Int study areas were a typical variety of *Passiflora biflora*, whereas the plants collected in the Wet study area were a subspecies or closely related species of *P. biflora* (with a nearly identical cyanogenic profile, referred to hereafter as *P. biflora*). One to four leaves were sampled and pooled per plant individual starting from the third leaf from the growing tip of the vine, and the samples were preserved fresh in 1ml 100% MeOH. In addition, 50 cm cuttings, all from different plant individuals, were sampled from the "Dry" ($n$ = 12) and "Wet" ($n$ = 12) study areas for greenhouse cultivation.

### Host plant greenhouse cultivation and treatments

Standard (std)-treated *P. biflora* were cultivated in greenhouses (75% relative humidity; 8-20 hrs: 25°C, light; 20-8 hrs: 20°C, dark) for use as oviposition plants for the parental generation and larval diet of the F1 families not included in feeding treatments. Std-plants were a greenhouse-cultivated stock of *P. biflora* established from plants originating from the Intermediate population in Panama. Vine cuttings were individually potted in a soil mixture (50% compost, 20% coir, 15% perlite, 15% sand or gravel), and watered three times weekly. The *P. biflora* cuttings collected from the Dry and Wet study areas were transported to the greenhouse and planted in pots with the same soil mixture as above. The watering treatments were initiated one month after potting when the plants had rooted and started new growth. Plants originating from the Dry and Wet study areas were divided equally into two watering treatments: the dry and wet treatments, leading to a total of four plant treatment groups. Plants in the wet treatment were watered three times weekly with soil remaining moist throughout the experiment. Additionally, wet-treated plants were placed near a hydrofogger (Hydrofogger 400 Simply Control), which increased relative humidity around the wet plants up to 100%. Plants in the dry treatment were kept at the standard 75% relative humidity and watered two to three times weekly with about 1/3 of the water volume compared to the wet treatment, such that the surface soil was allowed to dry and the leaves slightly droop between watering events. The dry treatment thus aimed to mimic conditions of drought stress. Exact volumes of water are not given, because the total amount of water was adjusted based on the size of the plant at the time of watering (as the vines gain mass, they require increasing amounts of water). All plants were given general plant fertilizer once weekly.

### Butterfly rearing and breeding design

Laboratory *H. erato* populations descendent of the wild Dry and Wet populations were established in greenhouse conditions (75% relative humidity; 8-20 hrs: 25°C, light; 20-8 hrs: 20°C, dark). Larvae were reared to adulthood on std-type *P. biflora*. All butterflies were marked with an individual identification number on the underside of the forewing. Butterflies were allowed to fly, mate, and oviposit freely in 2×2×2 m mesh cages with potted host plants (*P. biflora* std-type), and were fed with a standardized sugar solution diet *ad libitum* (20% sugar solution enriched with protein concentrate supplemented with amino acids (Vetark Professional Critical Care Formula; two scoops per 1 l of sugar solution). The sugar solution was served from artificial flowers and was replaced every second day. Dry and Wet-originating populations were kept in separate cages. The Wet population was unfortunately lost due to an infection. The Dry population maintained population sizes above several tens of individuals even at the peak of the infection, and therefore the experiment was continued with the Dry population. The butterfly cages were observed throughout the day for mating pairs. Mating individuals were identified based on their id numbers, and once a mating event was completed, the mated female was moved to an individual 2×2×2 m mesh cage with potted host plants *P. biflora* (std-type), and with the standardized sugar solution diet available *ad libitum*. Eggs of each female ($n$ = 14 mothers, of which 8 were mated with a known father, see Supplementary File 1) were collected twice weekly during the following two to three weeks following mating, and transferred into 60×35×35 cm mesh cages (with all full-sibling offspring of a female in one cage. Average no. of full-sibs/family = 8; Supplementary File 1) onto new growth harvested from std-type host plants. The vine cuttings were kept fresh by placing the end of each cutting through a hole in the lid of a water-filled container (which prevented the larvae from drowning). Fresh cuttings were added three times weekly, until all offspring had pupated. Six mated pairs (all with both mother and father known) and their offspring (average no. of full-sibs/family = 29, Supplementary File 1) were included in the feeding treatments, and the eggs of these six females were divided equally into four groups. Each group of eggs was transferred into separate 60×35×35 cm mesh cages onto new growth harvested from each of the four plant treatment groups. The cages were checked three times weekly for emerged adults, which were weighed and preserved in 100% MeOH (body excluding wings and one-half of thorax). All F1 individuals were sampled within three days following emergence, when they were still unmated and unfed.

**Butterfly toxicity analyses with $^1$H-NMR**

The concentrations of the two cyanogenic compounds biosynthesized by *Heliconius* larvae and adults, linamarin and lotaustralin, were analyzed from the butterfly samples using nuclear magnetic resonance ($^1$H-NMR). Sample extraction applied the procedure of Kim et al. (2010). The MeOH in which butterfly samples were preserved was evaporated and butterfly samples were dried by incubating at 45°C for 20 h. The dried samples were weighed (Denver Instrument SI-234, accuracy 0.1 mg) and homogenized in 2 ml Safe-Lock Eppendorph® tubes with an added sterile steel bead in an extraction/NMR solvent consisting of 400 µl

$KH_2PO_4$ buffer in $D_2O$ (pH 6.0) containing 0.1% (wt/wt) 3-(trimethylsilyl)propionic-2,2,3,3-d4 acid sodium salt (TSP) and 400 µl Methanol–$d_4$ ($CD_3OD$) 99.8%, using TissueLyser (Qiagen) at 30/s, 4*20 sec. Subsequently, samples were vortexed at maximum speed for 1 min, sonicated for 20 min (Eurosonic 22), and centrifuged 15 min at 13.2*$10^3$ rpm, before pipetting 625 µl of supernatant into a high-throughput NMR tube (Wilmad WG-1000-7 5*177,8 mm).

$^1$H-NMR was performed at the Finnish Biological NMR Center using Bruker Avance III HD NMR spectrometer (Bruker BioSpin, Germany) equipped with a cryogenic probe head and operated at $^1$H frequency of 850.4 MHz. Measurements were performed at 25°C. A standard cpmgpr1d pulse program (Bruker BioSpin; Carr and Purcell 1954) was chosen, using 32 scans, 32,768 data points, a spectral width of 10,204 Hz and a relaxation delay of 3 s. This pulse sequence is suitable for selective observation of small molecule components in solutions containing macromolecules. The data were processed and analyzed using Bruker TopSpin software (versions 3.2pl6 and 3.5pl6). The Free Induction Decays (FIDs) were multiplied by an exponential function equivalent to 0.3 Hz line-broadening factor before applying Fourier transformation. Transformed NMR spectra were corrected for phase distortions and calibrated according to the calibration compound TSP. The peaks of CNglcs compounds were identified based on comparison of sample $^1$H spectra with the $^1$H spectra of authentic reference samples of linamarin and lotaustralin. Concentrations of the CNglcs compounds were calculated based on the extracted areas of singlet peaks at 1.652 ppm (epilotaustralin) and 1.668 ppm (lotaustralin), and a duplet peak at 1.698 ppm (linamarin), quantified based on the extracted area of the calibration compound (TSP), and corrected for the sample dry mass. The values of CNglcs concentration in the wild-collected samples were normalized with a square-root-transformation before applying ANOVA to test for population differences in R 3.4.4 (R Core Team, 2018). Distributions of cyanogen traits were also explored by inspection of truncated weighted density distributions (accounting for the skewness of data towards near-zero values) using the “sm” R package (Bowman & Azzalini, 2014). For the analyses using common-garden data, untransformed values were used.

**Host plant toxicity and quality analyses**

The cyanogenic content of the host plant samples was analyzed using liquid chromatography-mass spectrometry (LC-MS/MS). A boiling method (Lai et al., 2015) was used for extraction, in which the leaf samples preserved in 1 ml 100% methanol were boiled for 5 min in a water bath. The samples were centrifuged at 10 000 g for 5 min, and the supernatant was filtered (Anapore 0.45 µm, Whatman), and diluted to 25%. Analytical LC-MS was carried out using an Agilent 1100 Series LC (Agilent Technologies, Germany) hyphenated to a Bruker HCT-Ultra ion trap mass spectrometer (Bruker Daltonics, Germany) following the procedure of Castro et al. (2019). Mass spectral data were analyzed with the native data analysis software (Compass DataAnalyses, Bruker Daltonics). CNglcs were identified by the molecular mass of their sodium adducts and by their fragmentation pattern (MS/MS). The amount of each compound was calculated based on extracted ion chromatogram (EIC) peak areas and quantified based on calibration curves of amygdalin (in the case of passibiflorin and passibiflorin diglycoside) and linamarin (in the case of tetraphyllin A/deidaclin). Quantification was corrected for spectrometer measurement efficiency and the sample dry mass, which was measured following the drying of extracted leaf samples at 45°C for 48 h.

We also measured overall host plant quality to estimate differences in nutritional values of the greenhouse-cultivated plant treatment groups, because larval-acquired nitrogenous resources have been proposed as an important source for cyanogen biosynthesis (Cardoso & Gilbert, 2013). Plant quality traits were assessed with Dualex® Scientific+ leaf clip meter (Cerovic et al., 2012), which measures chlorophyll content in µg/cm², epidermal flavonol content in absorbance units, nitrogen status (NBI: ratio between chlorophyll and flavonols; related to nitrogen/carbon allocation), and an anthocyanin index related to water-soluble pigments abundant in newly forming leaves and those undergoing senescence. The measurements were taken from the third leaf from the growth tip on the same day across all six plant individuals of each *P. biflora* treatment group ($n$ = 63 measurements on average per treatment group). ANOVA including plant origin, treatment and their interaction was applied in R 3.4.4 (R Core Team, 2018) to test for differences in CNglcs and plant quality traits between the treatment groups.

**Details on the estimation of variance components, heritability, evolvability and maternal effects in *Heliconius* biosynthesized cyanogenic toxicity**

When fixed effects are included in an animal model, the total random-effect variance is an underestimate of the total phenotypic variance ($V_P$), because it does not account for the part of the variance explained by fixed effects (Villemereuil et al., 2018). To overcome this, we computed the variance components attributed to the fixed effects of sex and feeding treatment using the method of Villemereuil et al. (2018). Here the fixed variance component ($V_F$) is calculated as $V_F = V(X\hat{b})$, where $X$ is the matrix of the values of cofactors, and $\hat{b}$ are the parameter estimates (here, the parameter estimates obtained from the fitted animal models). The standard errors of these fixed-effect variance components were calculated as the square root of the sampling variance of a variance (Lynch & Walsh, 1998). When calculating heritabilities we included the variance due to sex in the total phenotypic variance, but we did not include the variance due to the treatment because this component of the variance is unlikely to represent natural variation (Villemereuil et al., 2018).
